# Single cell profiling of functionally cured Chronic Hepatitis B patients reveals the emergence of activated innate and an altered adaptive immune response in the intra-hepatic environment

**DOI:** 10.1101/2022.04.26.489625

**Authors:** Balakrishnan Chakrapani Narmada, Atefeh Khakpoor, Niranjan Shirgaonkar, Sriram Narayanan, Pauline Poh Kim Aw, Malay Singh, Kok Haur Ong, Collins Oduor Owino, Jane Wei Ting Ng, Hui Chuing Yew, Nu Soibah Binte Mohamed Nasir, Veonice Bijin Au, Reina Sng, Nivashini Kaliaperumal, Htet Htet Toe Wai Khine, Hui Xin Ng, Su Li Chia, Cindy Xin Yi Seah, Myra HJ Alnawaz, Chris Lee Yoon Wai, Amy Yuh Ling Tay, Weimiao Yu, John Edward Connolly, Giridharan Periyasamy, Seng Gee Lim, Ramanuj Dasgupta

## Abstract

Hepatitis B surface antigen (HBsAg) loss or functional cure (FC), is considered the desirable therapeutic outcome for chronic hepatitis B (CHB) patients. However, the immuno-pathological biomarkers and underlying mechanisms remain unclear. Here we present a comprehensive single cell-transcriptomic atlas together with immune-phenotyping of disease-associated cell states (DACS) isolated from intra-hepatic tissue and matched PBMCs of either CHB or FC patients. We find that the intra-hepatic environment displays specific cell identities and molecular signatures that are distinct from PBMCs. FC is associated with emergence of an altered adaptive immune response marked by CD4 cytotoxic T lymphocytes (CD4-CTLs), and an activated innate response represented by liver-resident natural killer (LR-NK) cells. Overall, these findings provide novel insights into immuno-pathological cell states associated with FC that could serve as prognostic biomarkers.

## Introduction

Chronic Hepatitis B (CHB) virus infection affects more than 257 million people, mainly in the Asia-Pacific region leading to an estimated 800,000 deaths annually from cirrhosis and hepatocellular carcinoma (HCC) (1). The clearance of viral surface antigen (HBsAg) with or without appearance of anti-HBs antibodies, known as functional cure (FC), is now accepted as the desired therapeutic endpoint in the clinic (2). Current available antiviral therapies suppress viral replication by lifelong use but do not clear infection due to the persistent pool of covalently closed circular DNA (cccDNA) in hepatocytes (3, 4). The clearance of viral infection is largely dependent on a robust and long-lasting adaptive immune response. Virus-specific CD8 T cells are known to be involved in the clearance of viral infection in acute HBV-infected patients (5, 6). In CHB patients however, HBV-specific CD8 T cells are terminally exhausted due to chronic stimulation by high levels of HBV antigens (7). Recent studies have shown that impairment of adaptive immune response is also a function of the duration of exposure to viral antigens in addition to high antigen load (1, 6, 8). The specific function of the innate immune response in CHB remains debatable and relatively poorly understood. Innate immune response is generally thought to be functionally impaired in CHB (9–11). However, a recent study using a mouse model of HBV infection identified a specific Kupffer Cell (KC) population (constituting resident, embryonically-seeded macrophages in the liver), that was shown to reinvigorate the anti-viral function of HBV-specific CD8 T cells in the presence of IL2 (12). Additionally, other recent studies have revealed a positive correlation between increased cytotoxic activity of NK cells and liver inflammation, typically in patients who clear HBsAg upon cessation of Nucleoside Analog (NA) treatment (13). These studies allude to a potential function for innate immune cells in the resolution of CHB (11–13). Even though recent studies have explored the dynamics of the liver environment in different phases of CHB using multiplexed imaging and bulk RNA sequencing (14, 15) in biopsies, and fine-needle aspirates (FNA)-based single cell RNA-sequencing (scRNA-seq) (16), the interrogation of DACS in functionally cured patients has remained elusive. This presents a clear unmet need for a comprehensive, high-resolution investigation of key cell types or cell-states and regulatory cell-cell interactions that may modulate underlying mechanisms involved in sustenance of FC.

In this study, we sought to comprehensively characterize the cellular landscape associated with patients displaying sustained HBsAg loss (n=9) versus those with CHB (n=29), representing the two ends of the disease spectrum, by profiling a total of 91,502 single cells isolated from liver biopsies (n=47,109) and matched PBMCs (n=44,396). Through the systematic analyses of transcriptomic profiles, inferred cell-cell interactions, and flow cytometry-based validation studies, we present a comparative atlas that represents the single cell atlas of intra-hepatic environment and matched PBMCs of CHB patients versus those achieving FC. Our analysis identifies distinct populations of adaptive and innate immune cells, such as CD4-CTLs and NK cells, that may play essential function(s) in achieving and the maintenance of sustained HBsAg loss in FC patients. Altogether, our work provides deeper understanding of the disease-associated alterations in immune response while illuminating potential mechanistic drivers of FC. Our study also highlights that profiling the liver is indispensable for the discovery of tissue-specific DACS given our observation that alterations in the liver compartment are often not adequately reflected in matched PBMCs (17).

## Results

### Single cell atlas of liver and matched PBMCs from CHB and FC patients

Patients were categorized into CHB (n=29) where viral HBsAg is detectable, and FC (n=9) who displayed clearance of viral HBsAg and the sustained presence of anti-HBs antibodies (HBs-Ab) (Fig. 1A and Table 1). Total cells isolated from core needle biopsies of liver and patient-matched PBMCs were subjected to scRNA-seq using the 10x Chromium platform, and multiparametric flow cytometry (see Fig. 1A for overall workflow). For focused phenotypic analysis of immune cell subsets, CHB patients were stratified based on their quantitative HBsAg (qHBsAg), from high (>10000 IU/ml), medium (1000-10000 IU/ml), and low (0.05-1000 IU/ml), compared with FC patients, as shown in Table 2. Complete information on CHB disease phases based on EASL guidelines (2, 18), viral parameters, patient age, and ALT levels are provided in Tables 1 and 2.

**Figure 1.**
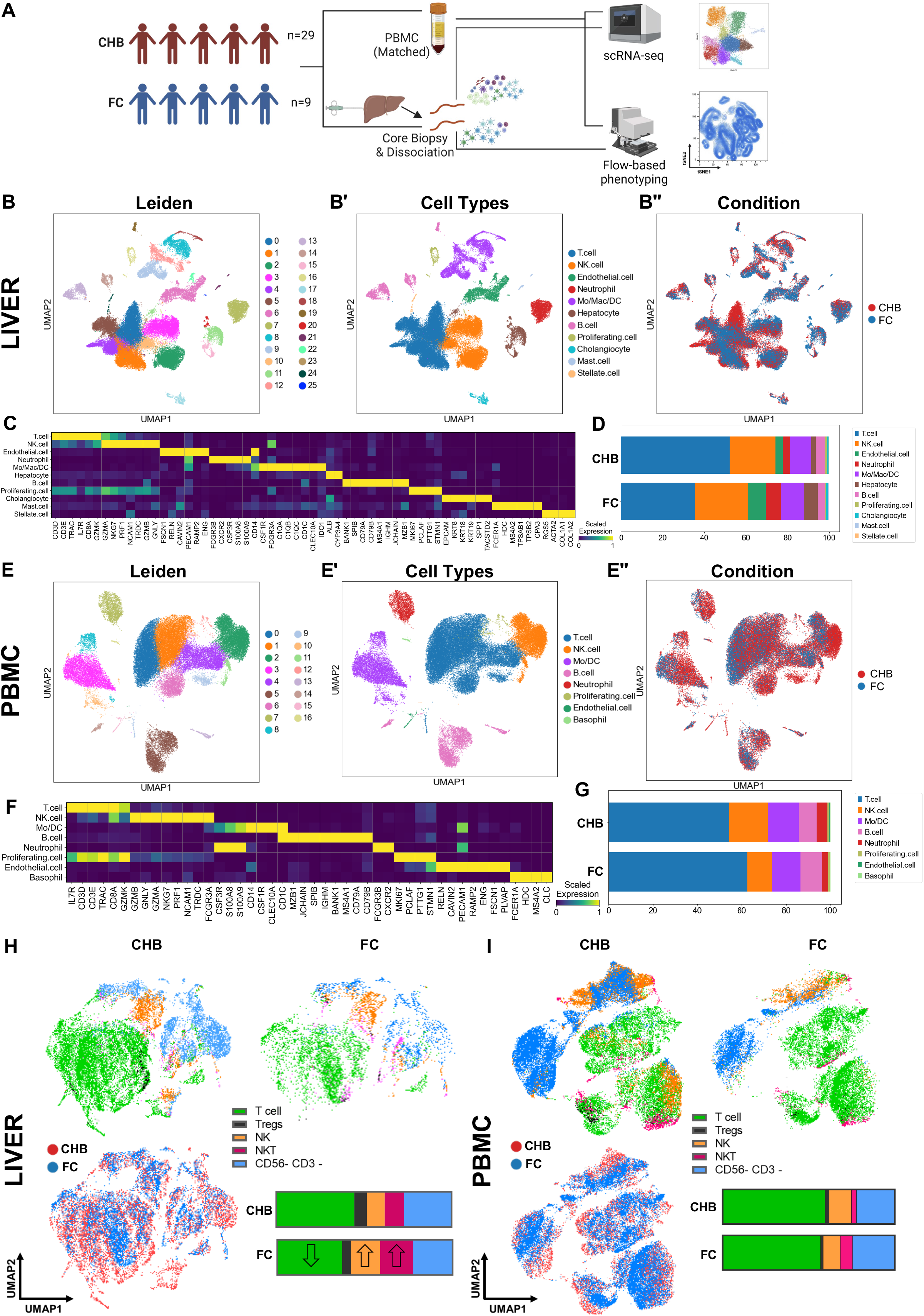
Single cell transcriptomic atlas of Liver and matched PBMCs isolated from CHB versus FC patients. **(A) Schematic of overall workflow** summarizing patient cohort, sample collection and experimental design. Matched PBMC and liver biopsy of CHB vs FC patients subjected to scRNA-seq, multiparameter flow-cytometry and multiplexed immuno-fluorescence. **(B)** UMAP projection of Leiden-based clustering of ~47,109 scRNA-seq libraries generated from the liver biopsies of CHB and FC patients identifies 26 distinct cell-states. **(B’)** Annotation of Leiden clusters based on the top differentially expressed genes (DEGs) and reference-based markers identifies 11 major cell types. **(B”)** Condition-wise annotation of scRNA-seq data from the two patient cohorts of CHB (red) vs FC (blue) within the liver. **(C)** Heatmap of cell type-specific markers within the liver compartment of CHB and FC patients. **(D)** Percentage-based proportionality bar graph representing cell types within the liver between CHB and FC patients. **(E)** UMAP projection of Leiden-based clustering of ~44,396 scRNA-seq libraries generated from PBMC of CHB and FC patients identifies 17 distinct cell-states. **(E’)** Annotation of Leiden-clusters based on the top differentially expressed genes (DEGs) and reference-based markers identifies 8 major cell types. **(E”)** Condition-wise annotation of scRNA-seq data from the two patient cohorts of CHB (red) and FC (blue) in PBMCs. **(F)** Heatmap of cell type-specific markers within PBMCs isolated from CHB and FC patients. **(G)** Percentage-based proportionality bar graph representing cell types in PBMCs from CHB or FC patients. **(H - I)** Multiparametric flow cytometry on isolated leukocytes from liver biopsies (H) and PBMC (I) from CHB or FC patients. UMAP projection of manually gated immune populations from liver or PBMC. Bar charts represent average relative frequency of the different immune subsets (including T cell, Tregs, NK, NKT as well as CD56-/CD3-cells). Note: down-arrows represent a decrease whereas up-arrows depict an increase in relative proportions of individual cell types between conditions (CHB or FC).

**Table 1:** Clinical and virological parameters of patients grouped based on the EASL guidelines.

| Characteristics | Patient group |  |  |  |  |
| --- | --- | --- | --- | --- | --- |
|  | HBeAg+chronic infection | HBeAg+chronic hepatitis B | HBeAg-chronic infection | HBeAg-chronic hepatitis B | FC |
| Number of patients | 0 | 10 | 7 | 12 | 9 |
| Age(years/mean) | NA | 37 | 53 | 55 | 54 |
| Sex (Male/Female) | NA | 6/4 | 3/4 | 6/4 | 2/6 |
| qHBsAg level (IU/ml) range (mean) | NA | 19768 | 1532.24 | 2940 | <0.05 |
| ALT level (IU/L) range(mean) | NA | 70.4 | 21.85 | 68 | 28 |
| HBV DNA level (IU/ml) range (mean) | NA | 1.06E+08 | 2.39E+04 | 2.03+06 | 0 |
| HBV genotype (A/B/C/D/E/ unknown) | NA | 1/3/6/0/0/0 | 0/4/2/0/0/1 | 0/4/4/1/0/0/3 | 0/1/5/0/0/3 |
| Naïve/NA/NA+IFN-a | NA | 10/0/0 | 4/3/0 | 8/3/1 | 2/1/6 |
| Anti HBs-Ab (IU/l) range (mean) | 0 | 0 | 0 | 0 | 93 |

**Table 2:** Clinical and virological parameters of patients. CHB patients are stratified based on qHBsAg level (IU/ml) (>10000, 1000-10000, 0.05-1000) and functional cure (FC).

| Characteristics | Patient group based on qHBsAg levels (IU/ml) |  |  |  |
| --- | --- | --- | --- | --- |
|  | >10000 | 1000-10000 | 0.05-1000 | Functional cure |
| Number of patients | 9 | 8 | 12 | 9 |
| Age(years/mean) | 43 | 46.2 | 55.4 | 54 |
| Sex (Male/Female) | 3/6 | 3/5 | 8/4 | 2/6 |
| ALT level (IU/L) range(mean) | 92.7 | 59.5 | 44.7 | 28 |
| HBV DNA level (IU/ml) range (mean) | 3.24E+08 | 1.26E+07 | 1.49E+04 | 0 |
| HBV genotype (A/B/C/D/E/ unknown) | 1/3/5/0/0/0 | 0/4/3/1/0/0 | 0/4/3/0/0/5 | 0/1/5/0/0/3 |
| HBeAg (Positive/Negative) | 8/1 | 2/6 | 0/12 | 0/9 |
| qHBsAg level (IU/ml) range (mean) | 4888 | 2989 | 186.3 | <0.05 |
| Naïve/NA/NA+IFN-a | 9/0/0 | 8/0/0 | 4/6/2 | 2/1/6 |
| Anti HBs-Ab (IU/l) range (mean) | 0 | 0 | 0 | 93 |

### Distinct alterations in the composition of adaptive and innate immune cells observed in liver versus matched PBMCs from FC patients

Single cell transcriptomic profiling of CHB and FC patients captured 47,109 cells within the liver comprising 26 distinct major cell types/states, as determined by Leiden-based clustering, representing epithelial (hepatocytes, and cholangiocytes), immune (T cells, B cells, NK cells, monocytes, macrophages, dendritic cells, neutrophils, and mast cells), endothelial and stellate cells (Fig. 1B-B’). Similarly, profiling of matched PBMCs revealed a capture of 44,396 cells constituting 17 different immune cell subtypes, mainly comprising T, NK, monocytes, DCs and B cells (Fig. 1E-E’). We employed Harmony-based integration (19) to eliminate variations arising due to batch differences and sequencing platforms and observed clear hierarchical clustering of each cell type and representation from each patient in the individual cell clusters. Cell type annotation of Leiden clusters was based on lineage-specific markers adapted from earlier scRNA-seq studies profiling the liver (20–22) (Fig. 1C and 1F), such as T cells (*CD3E, CD3D, IL7R, CD8A, CD8B, TRAC*), NK cells (*NCAM1, GNLY, NKG7, TRDC, PRF1*), myeloid cells (*CD14, CSF1R, FCGR3A, C1QA, C1QB, IDO1, CLEC10A*), B cells (*MS4A1, MZB1, CD79A, JCHAIN*), stellate cells (*ACTA2, RGS5, COL1A1*), endothelial cells (*PECAM1, RELN, ENG, FSCN1, CAVIN2*), and hepatocytes (*ALB, CYP3A4*). Importantly, we successfully captured an adequate representation of neutrophils (*FCGR3B, CSF3R, S100A8, S100A9*) by slight modification of single cell dissociation and sequencing methodologies. This allowed us to generate a comprehensive liver profile and delineate changes associated with FC, compared to CHB patients (Fig. 1B”, D, E” and G).

Initial analysis of broad cell types revealed that compared to CHB, the FC patients display a marked reduction of the T cells along with a concomitant increase in NK cells and neutrophils in the intra-hepatic environment (Fig. 1D). Intriguingly, we failed to observe similar trends in the peripheral circulation upon analysis of PBMCs from matched patients (Fig. 1G). Notably, multiparametric flow cytometry (Fig. 1H) also confirmed the reduction in T cells and an increase in NK/NKT population in the liver biopsies of FC patients, while the peripheral circulation failed to capture similar changes (Fig. 1I). Overall, these observations suggest that there may be specific alterations in immune cell composition within the livers of functionally cured patients that may not always be reflected accurately in PBMCs.

### FC patients exhibit marked loss of exhaustion markers in adaptive immune cells with a concomitant gain in innate immune response

Next, we sought to determine the phenotypic changes in T and NK cell populations among FC and CHB patients. First, we subdivided the major T and NK clusters (identified in Fig. 1B, 1B”) followed by re-clustering of these cell types to reveal heterogeneity in their phenotypic states and investigate how they are altered upon functional cure compared to CHB (Fig. 2A-B’). Deep phenotyping of the sub-clustered T and NK cells using cluster-specific differentially expressed genes (DEGs) from the scRNA-seq data resulted in the identification of distinct cellular subtypes or cell-states including CD4 T helper cells (*CD4, IL7R, CCR7*), CD4 Cytotoxic T lymphocytes or CD4-CTLs (*IL7R, KLRG1, PRF1, TNF*), CD8 T effector cells (*CD8A, KLRG1, PRF1, KLRK1, IFNG*), exhausted CD8 T cells (*CD8A, LAG3, FASLG, PDCD1, HAVCR2, TIGIT^high^*), memory-like CD8 T cells (*CD8A, IL7R, TNF, IFNG, TNFSF10^high^*), Tregs (*FOXP3, IL2RA, CTLA4^high^, BATF^high^, TIGIT*), Liver Resident NK or LR-NK (*CXCR6, EOMES, NCAM1^high^*), conventional NK or cNK (*NCAM1^low^, FCGR3A, GNLY, GZMB, PRF1^high^*), and NKT cells (*CD3D, NCAM^low^, GZMK, GNLY, PRF1*) (Figs. 2A, B, G and Supp. Fig. S1). In particular, we noted a marked reduction in the exhausted PDCD1+ CD8 T cells as well as an almost complete loss of FOXP3+ Tregs in FC patients (Fig. 2A, A” and 2B, B”). The exhaustion markers such as TIGIT, LAG3 and PDCD1 are significantly reduced in FC patients (as depicted in Fig. 2A’, 2B’, and violin plots in Fig. 2C-D). However, these changes are mainly reflected in the liver (Fig. 2C), rather than in PBMCs, likely due to the lower frequency of these cells in the peripheral circulation (Fig. 2D). Similarly, the decline in the frequency of FOXP3+ Tregs in FC patients compared to CHB was more evident in the liver, compared to matched PBMC samples (Fig. 2A, 2A’, and 2B’). Importantly, flow cytometry analysis further corroborated these findings as shown in Fig. 1H and 1I. Direct *ex vivo* flow cytometry analysis on global CD8 and CD4 T cells in liver and PBMCs also revealed a markedly lower frequency of PD1+ exhausted T cells in FC patients. This was evident not only in the CD69+ tissue resident CD8/CD4 T cells, but also in the transiting CD69-T cells. Stratifying the CHB patients based on the levels of qHBsAg showed that the reduction of exhausted CD4 and CD8 T cells correlate with reduction of qHBsAg (Fig. 2E).

**Figure 2.**
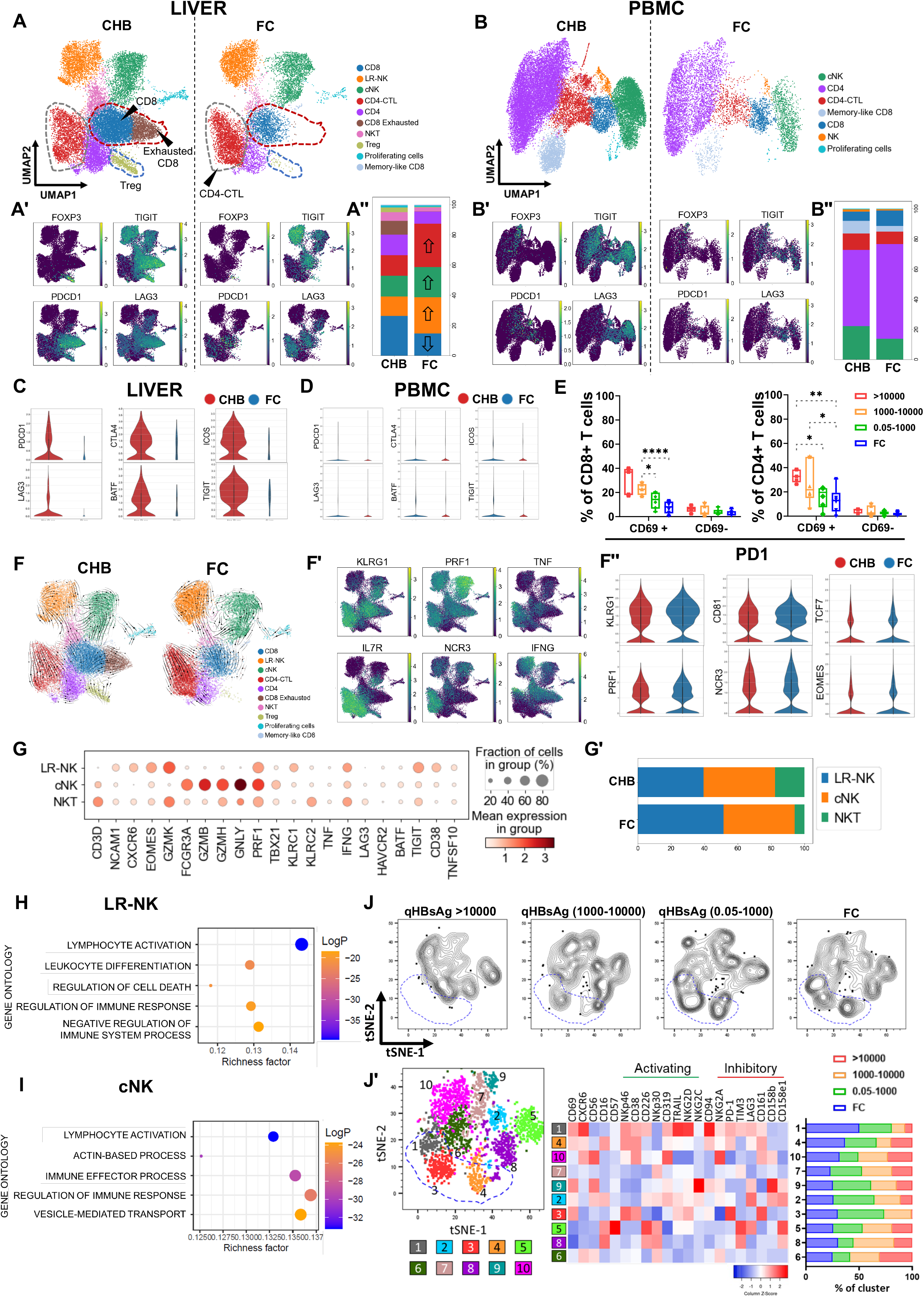
Patients exhibiting FC display loss of exhaustion markers, a gain of innate and an altered adaptive immune response. **(A-B”)** Comprehensive transcriptomic analysis of NK and T cells identified from CHB and FC patients, either in liver (A-A”) or matched PBMCs (B-B”). **(A)** Loss of exhausted CD8 T cells (brown cluster marked within red dashed line) and Tregs (marked by blue dashed line) in FC patients. Note the marked reduction in exhaustion markers *TIGIT, PDCD1, and LAG3* amongst the exhausted CD8 T cells and the expression of FOXP3+ Tregs (A’). Percentage-based proportionality bar graph summarizes the differences in composition of T cell subsets between CHB and FC patients (A”). Note: down-arrows represent a decrease whereas up-arrows depict an increase in relative proportions of individual cell types between conditions (CHB or FC). **(B-B”)** Similar analyses in PBMCs of CHB and FC patients reveal only a modest trend in the loss of exhausted CD8 T cells and Tregs. **(C-D)** Violin plots showing expression of specific exhaustion markers *PDCD1, CTLA4, ICOS, LAG3, BATF, and TIGIT* between CHB and FC patients in liver (C) and PBMC (D). **(E)** Frequency of PD1+CD8 and CD4 T cells shown in tissue resident (CD69+) and transiting (CD69-) populations in liver (CHB patients stratified based qHBsAg levels). **(F)** Steady-state RNA velocity of NK and T cell clusters in CHB and FC patients within liver compartment. Note: arrows depict the directionality of cell-state transitions. **(F’)** Increased expression of specific effector cell markers on CD4-CTL clusters (*KLRG1, PRF1, TNF, IL7R, NCR3, IFNG*) (“red” cluster within grey dashed line in panel A) **(F”)** Violin plots for differential expression of markers related to cytolytic function of CD4-CTLs (*KLRG1, CD81, TCF7, PRF1, NCR3, and EOMES*) between CHB and FC patients. **(G-G’)** Dot plot and percentage-based proportionality chart depicting differential expression and frequencies of distinct NK cell subsets (LR-NK, cNK and NKT) within liver compartment between CHB and FC patients. **(H-I)** Gene ontology (GO) pathway analyses on LR-NK and cNK cells from CHB and FC patients. **(J-J’)** Multiparametric flow cytometry analysis on total NK cells isolated from liver. tSNE clustering (J) and PhenoGraph analysis (J’) on CHB patients (stratified based qHBsAg levels) and FC patients (J’). Z score normalized expression of activating and inhibitory receptors in each cluster shown in heatmap (J’). Bar chart depicts frequency of each cluster in CHB patients (stratified based on qHBsAg levels) and FC patients. Note: qHBsAg stratification based on the following parameter: IU/ml >10000; 1000-10000; and 0.05-1000. Statistics in panel E calculated using two-way ANOVA with Tukey test, *P≤ 0.05, **P≤ 0.01, ****P ≤0.0001.

Most unexpectedly, concomitant with the loss of exhausted CD8 T cells and Tregs, we observed a distinct increase in a population of CD4 T cells that displayed markers typically associated with effector cytotoxic lymphocytes (*PRF1, GZMA, TNF, NCR3, IFNG* (Supp. Fig. S1)), hence referred to as CD4-CTLs, in patients displaying FC (Fig. 2F’, F”). Trajectory analysis via RNA velocity (23) (Fig. 2F) suggested that the emergence and increase of CD4-CTLs in FC patients is most likely a result of naïve CD4 cells differentiating into CD4-CTLs, possibly under the stimulation of IL2-STAT5 signaling and IFNγ, as inferred by pathway analysis (data not shown). Notably, the trajectory analysis also revealed a clear demarcation between CD4-CTL and CD8 T cells even though they share common marker genes like *CD8* and *IL7R*, suggesting a distinct cell-of-origin for CD4-CTLs, which may subsequently engage in an altered adaptive immune response. Furthermore, deep phenotyping revealed that CD4-CTLs express markers associated with terminally differentiated, antigen-experienced CD4 T cells (*IL7R, KLRG1*) that display unique characteristics of inflammatory, cytotoxic effector cells including *TNF* (TNFα) and *IFNG*. Interestingly, they also express the *NCR3* gene resembling features of innate, NK-like cells (Fig. 2F’ and as described in (24)), suggesting a gain of innate-like function in CD4-CTLs. Additionally, we observed an increase in *KLRG1* and *CD81* in FC patients indicating antigen exposure of these cells (25) compared to other cluster-defining marker genes (*PRF1, NCR3, TCF7, TNF, IFNG, EOMES*) that remain unchanged between CHB and FC patients (Fig. 2F”). Altogether, these observations underscore a shift in the intra-hepatic environment from an immune-suppressed to an immune-activated state in functionally cured patients.

Trajectory analysis also exposed remarkable plasticity of the CD8 T cells (Fig. 2F). We identified two populations of CD8 T cells; one that is observed predominantly in CHB (right segment of CD8 cluster), which also gives rise to the exhausted CD8 T cells, marked by the expression of *PDCD1, LAG3 and TIGIT*(Fig. 2F). FC was marked by a striking disappearance of the exhausted CD8 T cell cluster. Intriguingly, the remaining subset of CD8 T cells revealed trajectories that suggest their transition into NKT and cNK cells. The cNK and NKT clusters in turn appear to contribute towards the LR-NK cells that are increased in FC patients suggesting the transition from CD8 to NKT or cNK to LR-NK might promote an innate immune active state. Altogether, this data suggests the presence of an altered adaptive immune response driven by CD4-CTLs expressing effector proteins (*PRF1, TNF, IFNG*) and pathways such as IL2, STAT5; and also highlights the plasticity of T cells to acquire innate-like features and likely, function (26–29).

In addition to the increase in CD4-CTLs, FC patients also displayed an increase in the classical innate immune response. Analysis of liver scRNA-seq data showed a distinct increase in the LR-NK cells (*CXCR6, EOMES*) (Fig. 2G and G’), which have been suggested to perform a more immune regulatory role by secreting cytokines and engaging with signaling interactions with various cell types, such as recruiting DCs to the site of inflammation and promote cross presentation of antigens to remodel the liver environment (30–33). The *FCGR3A+* conventional NK (cNK) cells, which are directly involved in antibody dependent cellular cytotoxicity (32) and known to promote anti-viral T cell response (33), also show a mild increase in FC patients (Fig. 2G, and G’). As shown in Fig. 2H and I, pathway analysis on gene expression signatures strongly suggested an increase in immune effector function in both cell types with specificity towards cytokine-chemokine secretion in LR-NK and cNK cells. Furthermore, direct *ex vivo* flow cytometry analysis in liver revealed a gradual enrichment of CXCR6+ CD69+ NK clusters with declining levels of qHBsAg (Fig. 2J). These cells displayed elevated expression of activating receptors (NKp46, CD38, NKG2D and TRAIL) (Fig. 2J’) and lower expression of inhibitory receptors, such as PD1 and TIM3 in FC patients. We suggest that the CXCR6+ CD69+ NK cells are representative of the LR-NK cells identified in the scRNA-seq analysis and have been described in earlier studies (30, 31). That said, some of the LR-NK cells in FC patients also displayed moderate-to-high expression of other inhibitory receptors, such as PD1 and LAG3 (Fig. 2J’ cl1,3,4). Based on these markers we identified CD69+ CXCR6^high^ CD16- (Fig. 2J’ cl 1) and CD69+ CXCR6^low^ CD16+ (Fig. 2J’ cl 4) populations that emerged with declining HBsAg levels and were enriched in FC. These populations are characteristic of the LR-NK and cNK respectively, as identified in scRNA-seq data (Fig. 2G). Interestingly we also identified another population marked by CD69+ CXCR6-CD16- that was enriched with low qHBsAg levels (Fig. 2J’ cl 3). Altogether, multiparametric flow cytometry data suggests that the effector cytotoxic function of tissue-resident NK cells must be carefully moderated by the balanced expression of activation and inhibitory receptors. These observations suggest a gradual but inverse trend of increased activated NK cell function with decreasing qHBsAg levels, which highlights a biological gradient of enhanced innate immune function associated with FC (34).

### Loss of senescence and gain of proliferation capacity in global T cells correlates with reduced qHBsAg

We utilized flow cytometry to comprehensively characterize the phenotype of global CD8 and CD4 T cell populations found within the liver and PBMCs of CHB patients versus those exhibiting FC (Fig. 3A-D). Specifically, we explored whether the phenotypic changes in CD4 or CD8 T cell populations were correlated with qHBsAg level. As depicted in Fig. 3A and C, tissue resident CD4 and CD8 T cells, marked by CD69+ population, showed a significantly higher frequency of memory-like T cells (CD127+ CD8 and CD127+ CD4 T cells) that correlated with a reduction of HBsAg level and in FC patients. This was also associated with a lower frequency of terminally differentiated T cells (HLA-DR+ T cells and CD57+ T cells) and increased frequency of CD28+ CD4 or CD8 T cells (35). Peripheral CD4 T cells showed somewhat similar phenotypic changes to those found in the liver compartment. In PBMCs, frequency of terminally differentiated CD4 T cells was slightly reduced in FC patients whereas we observed an increase in memory-like T cells (Fig. 3B). CD8 T cells in PBMCs however did not capture similar phenotypic changes as those found in the liver (Fig. 3D). Surprisingly, the frequency of CD57+ CD8 T cells increased in FC patients while the frequency of CD28+ CD8 T cells remained lower compared to CHB (Fig. 3D). Once again, these observations revealed distinct differences not only in the robustness of the immuno-phenotypic changes, but also in the patterns observed between the two tissue types (liver vs PBMCs). Loss of exhaustion and senescence combined with an increase in memory-like T cells suggests a gain in effector phenotypes and proliferative capacity in CD4 and CD8 T cells in the absence of viral antigens in FC patients (36, 37). Incidentally, increase in frequency of memory-like T cells has also been observed in HBV-specific T cells in patients with resolved acute HBV infection (36), further supporting the notion that similar mechanisms may drive viral clearance in CHB patients upon treatment.

**Figure 3.**
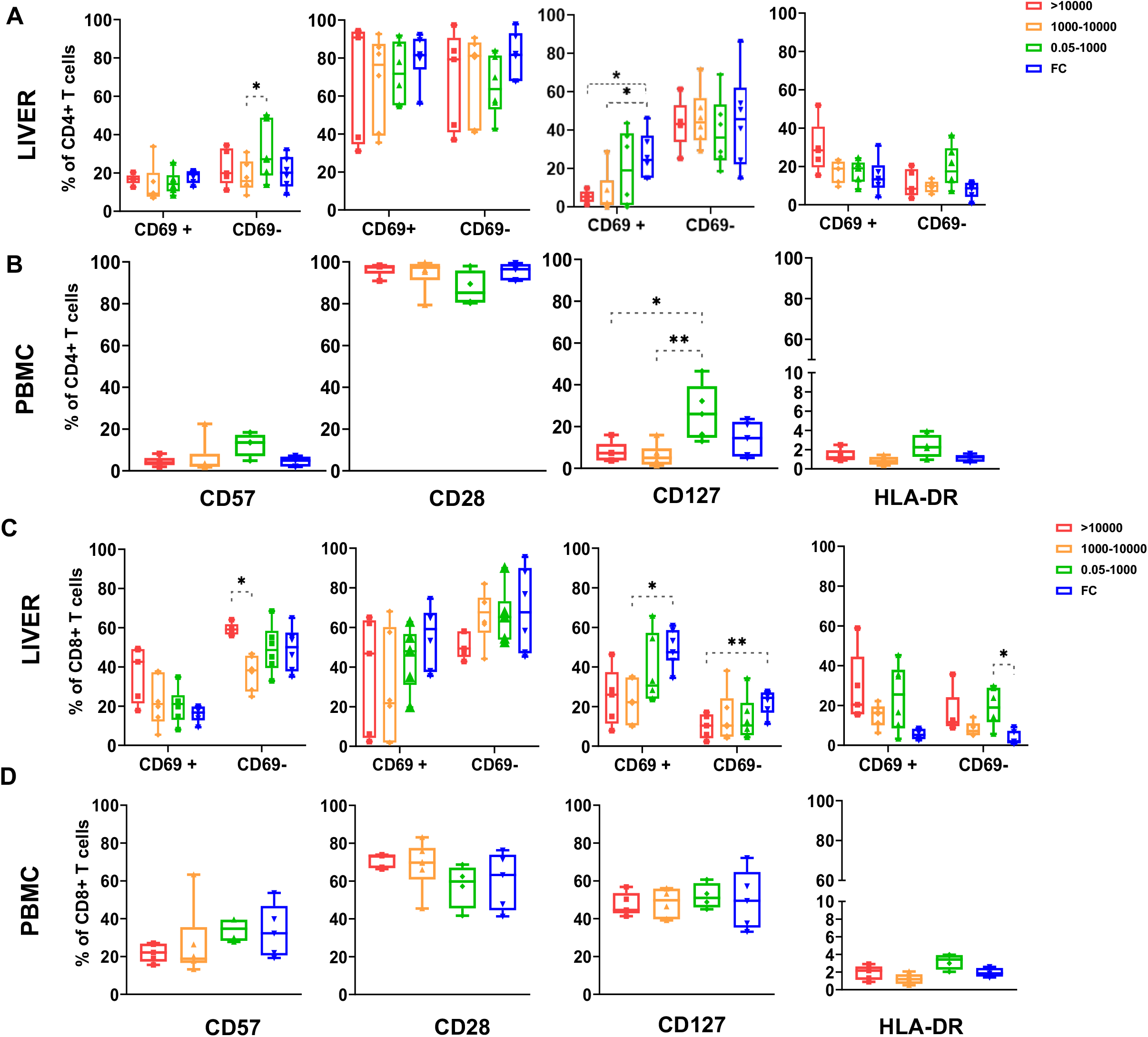
Phenotypic analysis of total T cells in patients exhibiting FC reveal emergence of effector T cells as key regulators of viral clearance. **(A-D)** Deep phenotyping of global CD8 and CD4 T cells in liver (CD69+ and CD69-) and PBMC of patients using multiparametric flow cytometry analysis (grouped based on qHBsAg). The frequencies of CD4 T cells expressing CD57, CD28, CD127 and HLA-DR in liver (A) and PBMC (B). Frequencies of global CD8 T cells positive for CD57, CD28, CD127 and HLA-DR in liver (C) and PBMC (D). Note: qHBsAg stratification based on the following parameter: IU/ml >10000; 1000-10000; and 0.05-1000. Statistics in panel E calculated using two-way ANOVA with Tukey test, *P≤ 0.05, **P≤ 0.01.

## Discussion

There remains a significant knowledge gap in our understanding of the underlying immuno-pathological events, especially within the intra-hepatic tissue, in CHB patients who have achieved functional cure. This is due both to the relatively low percentage of functionally cured patients (whether treatment-naïve or under lifelong antiviral therapy), as well as the limited access to their liver tissue. Moreover, the distinctive features of the liver cannot be completely captured by profiling the readily accessible PBMC samples, which further widens this knowledge gap (4, 18, 38). A systematic investigation is warranted to identify not only distinct immuno-pathological cell states that are relevant to the disease, but also potential signatures that correlate with disease resolution, and eventually FC. Despite intra- and inter-patient heterogeneity (39), through the comprehensive single cell profiling of both liver and matched PBMCs isolated from CHB vs FC patients, we identified both convergent and divergent gene expression-based DACS as well as their putative functions that may be reflective of mechanisms and biomarkers of FC.

Even though our dataset has been generated from a spectrum of patient samples from different phases of CHB and from functional cure, we identified two common and important denominators. First, there are profound differences in the DACS and molecular signatures identified between the liver and peripheral circulation. While there is convergence in the global T and NK cell response (Fig. 1, 2), emergence of pro-inflammatory innate immune response (neutrophils), marked loss of exhausted T cells and Tregs, and increased frequencies of specific primed NK subtypes (Fig. 2) were only detectable in the liver, thereby making intra-hepatic studies indispensable for the comprehensive identification of DACS. Recent studies also underscore the importance of interrogating the liver environment to identify distinct DACS specific to the liver environment that may otherwise be missed by exclusively analyzing PBMCs (16, 38, 40–42). Second, we identified a shift in the immunological profile of the liver from an immune-suppressed environment found in CHB patients to a pro-inflammatory, immune-active environment in FC patients. Such remodeling could occur due to the increased frequency of LR-NK cells with high expression of activation receptors, such as CD38, TRAIL, NKp46 and NKG2D and moderate levels of inhibitory receptors, such as PD1 and TIM3 (Fig. 2G, and cl1 in Fig. 2J’), long after the loss of HBsAg. Therefore, we hypothesize that these NK cells are primed, with memory-like features (43–46) that retain their capacity to elicit a coherent immune response when re-exposed to the virus, or to ensure long-term viral suppression during FC (29, 44, 47–49). In addition, the emergence of an altered adaptive immune response steered by CD4-CTLs (*PRF1, GZMA*) (Fig. 2), may also contribute towards the immune-active state in FC patients. Together, the data leads us to suggest that low-grade inflammation may be required for the maintenance of the “functionally cured” state.

Findings from this study lays the foundation for the functional interrogation of the key immunological events during the process of HBsAg loss. However, the loss of exhausted T cells, gain of innate immune activity, and the emergence of effector CD4-CTLs during the transition towards viral antigen clearance may serve as novel cell state-based prognostic biomarkers that can guide clinical management of CHB. In addition, findings from our studies may also suggest a therapeutic window of opportunity in patients that display reduced levels of viral antigens and exhaustion markers for the application of next-generation host-directed therapies, such as therapeutic vaccines, checkpoint inhibitiors (50) and immune modulators that can activate the innate and/or an altered adaptive immune response to facilitate the achievement of FC.

## Online Methods

### Materials and methods

#### Study approval

Peripheral Blood and biopsy sampling as well as clinical assessment were performed at the National University Hospital (Singapore) in accordance with the Declaration of Helsinki. Written informed consent was obtained. The study was reviewed and approved by Singapore National Health Group Domain Specific Review Board (DSRB reference number 2000/00828).

#### Patients

Paired blood and biopsy of 29 Patients with chronic Hepatitis B infection (across different phases of CHB based on EASL guidelines) (2, 18) and 9 patients who have achieved functional cure (qHBsAg <0.05) were obtained for this study. Two needle core biopsies were obtained from patients with signed consent using the biopince liver biopsy instrument. Tables 1 and 2 summarize the clinical and virological parameters of all patients.

#### Clinical and virological parameters

HBV DNA levels were measured in all patients (COBAS AmpliPrep/COBAS TaqMan HBV test v2.0; Roche Molecular Diagnostics) and viral parameters (qHBsAg, HBeAg, and anti-HBe levels)were assessed with Roche COBAS 8000 (e801) system using Electrochemiluminescence Immunoassay (ECLIA) method, and HBV genotype was recorded where available. All patients included were HCV, HDV and HIV negative.

#### Single cell dissociation for scRNA-Seq

Liver biopsies and PBMC were collected from patients with institutional ethics approval as described earlier. Liver biopsies were processed based on an enzymatic dissociation protocol adapted from Sharma et al. 2020 (21). Briefly, biopsies were stored and transported in HypoThermosol® FRS Preservation Solution (Sigma H4416-100ml) solution at 4°C until further processing. The tissue sample was first minced to 2-3mm pieces using sterile scalpels. Collagenase P solution (from Roche (11249002001) 2mg/ml in DMEM) was added to the tissue and the tissue was kept in a 37°C shaking incubator to allow single cell dissociation for 10-15min. Cell suspension was washed with PBS (no Mg2+ or Ca2+) with 1% w/v BSA, and 1mM EDTA solution filtered through 70 μm cell strainer to remove cell debris. The cells were then centrifuged at differential speeds to collect parenchymal fraction (100xg) and non-parenchymal fractions (400xg). If red blood cells were seen in the pellet, RBC lysis was performed using 10x RBC lysis buffer (Biolegend 420301) for 5min before proceeding to the next wash. Filtered through 40μm filter and washed with PBS (no Mg^2+^ or Ca^2+^) with 1% w/v BSA solution. Cell count and viability of the dissociated cells was evaluated by Trypan Blue staining before combining the parenchymal and non-parenchymal fractions in a 1:1 ratio to make up 30μl of 1000cells/μl processing for 10x 3’ Single cell RNA seq v3 preparation.

#### Flow cytometry sample preparation

To isolate the leukocytes, liver biopsy was subjected to mechanical disruption and filtration through a 70μm cell strainer (BD biosciences). Removal of the parenchymal cells and isolation of lymphocytes were done through centrifugation at 50xg without brake. Isolated lymphocytes were washed, counted to evaluate the cell viability and used immediately after isolation for further flow cytometry related analysis.

PBMC isolation from heparinized blood was carried out by density centrifugation with Ficoll-Hyperpaque Plus (GE Healthcare). Isolated PBMCs were cryopreserved in 10% dimethyl sulfoxide (DMSO) (Sigma Aldrich) prior to further use (51).

#### Single cell RNA seq sample preparation, cDNA & library generation, sequencing

An aliquot of the prepared single cells was counted, and cell concentration was adjusted to 800 cells/ul. GEM generation and 3’ mRNA-seq gene expression libraries were prepared using the Chromium Next GEM Single Cell 3’ GEM, Library and Gel Bead Kit v3.1 (10x Genomics) according to the manufacturer’s guidelines. Briefly, 16.5ul of cells were added into the master mix for a targeted cell recovery of 8000 cells. This was then loaded into a single-cell chip which was then inserted into the 10x Genomic Chromium controller for GEM generation. Once the run was completed, the samples were carefully transferred into clean tubes for reverse transcription. The samples were then purified using Dynabeads followed by 12 cycles of cDNA amplification. After that, samples were purified using SPRI-select reagent before profiling them on the Bioanalyzer (Agilent). Next, genomic scRNA libraries were constructed. This consisted of a series of steps involving fragmentation, end repair, A-tailing, adaptor ligation and a final amplification of 14 cycles. Finally, using SPRIselect reagent, the samples were purified and the required size of 300-700 bp was selected. Quantification and quality check was done using Bioanalyser (Agilent). Each of the samples XHL102 to XHL135 were sequenced in-house on a single lane on HiSeq 4000, whereas samples XHL232 to XHL343 were pooled with maximum of 8 samples in each lane and sequenced on NovaSeq S4 (Novogene). Both platforms generated paired-end reads with a length of 151 bp.

#### Single cell RNA-seq Analysis

Single-cell datasets generated in-house from CHB and FC patient livers and PBMC samples were processed using the Cellranger 6.0.1 pipeline and were mapped to the human reference genome (GrCh38). The output, raw feature-barcode matrix, was then imported into ScanPy (v1.7.1). Cells of low quality (cells with less than 200 genes per cell) and cells expressing rare genes (genes expressed by less than 30 cells) were then removed from the matrix. Cells with a mitochondrial percentage greater than 20% or cells with the number of detected genes (NODG) less than 200 were excluded from the matrix. An upper NODG limit of 6000 genes was used and the putative cell doublets were removed using the Scrublet package (52). The cell counts were then normalized using *scanpy.pp.normalize_per_cell* with a scaling factor of 10,000 and gene expression was scaled to unit variance and a mean value of 0 using *scanpy.pp.scale*. Dimensionality reduction of the data was done with PCA using the *scanpy.tl.pca* function and the *scanpy.pl.pca_variance_natio* plot was used to determine the inflection point after which no remarkable change in the variance was observed. The neighborhood graph for clustering was calculated using *scanpy.pp.neighbors* while the *scanpy.tl.leiden* function was used to cluster the cells using Leiden clustering. Batch effects were observed after clustering which could be attributed to the samples being sequenced using two different platforms, NovaSeq and Illumina HiSeq 4000. The batch correction pipeline, Harmony was used to integrate the samples together. All steps starting from dimensionality reduction were repeated until well-defined leiden clusters were obtained. Differentially expressed genes across the leiden clusters were determined using *scanpy.tl.rank_genes_groups* and were used to check cluster validity. Cell subtypes were then assigned to these clusters using annotation pipelines like singleR and Azimuth (53). Manual annotation using well-known cell type markers from available literature were also employed for confirmation. Each individual cell subtype was segregated and undesirable sources of variations like variable library sizes, mitochondrial genes, cell cycle genes were regressed out using the *scanpy.pp.regress_out* function. Cell subtype were then re-clustered and the resulting leiden clusters specific to each cell type were analyzed between CHB and FC patients using RNA Velocity (23) for specific cell type trajectory analysis.

#### Pathway analysis

Differentially expressed genes between clusters or between the two conditions (CHB and FC) were determined through Wilcoxon statistical tests (p-value<0.01). Upregulated pathway analysis of these DEGs was performed using open-source application Metascape (54) and visualized through the ggplot2 function in R.

#### Direct *ex-vivo* flow cytometry analysis

Freshly isolated leukocytes from liver biopsies and thawed PBMCs were stained with Live/Dead cell viability stain (Invitrogen, California, USA) followed by surface staining for 30 minutes on ice. All antibodies used for direct *ex vivo* analysis of T and NK cells are summarized in Tables 3 and 4. Cells were acquired on BD FACSymphony™ A5 (BD Biosciences) cell analyzer for NK and T cell analysis. Data was analysed using FlowJo version 10 (BD Biosciences). Heatmaps were generated using Heatmapper (55).

**Table 3:** List of antibodies used for multiparametric flow cytometry analysis of NK cells.

| Fluorochrome | Antibody | Catalog no | Clone | Source | Dilution |
| --- | --- | --- | --- | --- | --- |
| BUV395 | CD226 (DNAM-1) | 742498 | DX11 | BD Bioscience | 25x |
| BUV496 | CD45 | 750179 | HI30 | BD Bioscience | 25x |
| BUV563 | CD337 (NKp30) | 750179 | P30-15 | BD Bioscience | 25x |
| BUV615 | CD57 | Custom | NK-1 | BD Bioscience | 200x |
| BUV661 | CD335 (NKp46) | Custom | 9E2/Nkp46 | BD Bioscience | 100x |
| BUV737 | CD279 (PD1) | 612791 | EH12.1 | BD Bioscience | 25x |
| BUV805 | CD16 | Custom | 3G8 | BD Bioscience | 200x |
| BV421 | CD253 (TRAIL) | 564243 | RIK-2 | BD Bioscience | 25x |
| BV480 | CD186 (CXCR6) | Custom | 13B 1E5 | BD Bioscience | 25x |
| BV570 | CD3 | Custom | UCHT1 | BD Bioscience | 100x |
| BV605 | CD158b<br>(KIR2DL2/L3/S2) | 743453 | CH-L | BD Bioscience | 50x |
| BV650 | CD336 (NKp44) | 744302 | p44-8 | BD Bioscience | 25x |
| BV711 |  |  |  |  |  |
| BV750 | CD366 (TIM3) | Custom | 7D3 | BD Bioscience | 100x |
| BV786 | CD158e1 (KIR3DL1) | 555967 | DX9 | BD Bioscience | 100x |
| BB515 | CCR5 | 565153 | DX26 | BD Bioscience | 100x |
| BB630 | CD94 | Custom | HP-3D9 | BD Bioscience | 100x |
| BB660 | CD69 | Custom | FN50 | BD Bioscience | 200x |
| PerCP-Cy5.5 | CD314 (NKG2D) | 320818 | 1D11 | Biolegend | 25x |
| BB790 | CD161 | Custom | DX12 | BD Bioscience | 25x |
| PE | CD159c (NKG2C) | R&D<br>FAB138P | 134591 | R&D Systems | 25x |
| PE-CF594 | CD56 | 564849 | NCAM16.2 | BD Bioscience | 200x |
| PE-Cy5 | CD38 | 561823 | HIT2 | BD Bioscience | 50x |
| PE-Cy7 | CD223 (LAG3) | 369310 | 11C3C65 | Biolegend | 25x |
| AF647 | CD319 (CRACC) | 564338 | 235614 | BD Bioscience | 10x |
| AF700 | CD159a (NKG2A) | R&D<br>FAB1059N | 131411 | R&D Systems | 25x |
| APC-H7 | CD14 | 561384 | M5E2 | BD Bioscience | 50x |
| APC-H7 | CD19 | 560727 | HIB19 | BD Bioscience | 25x |

**Table 4:** List of antibodies used for multiparametric flow cytometry analysis of T cells.

| Fluorochrome | Antigen | Catalog no | Clone | Source | Dilution |
| --- | --- | --- | --- | --- | --- |
| **FITC** | KLRG1 | 367714 | SA231A2 | Biolegend | 200x |
| **PE** | NKG2D | 557940 | 1D11 | BD Bioscience | 50x |
| **BB660** | CD69 | 624295 | FN50 | BD Bioscience | 200x |
| **BB700** | CD56 | 566400 | B159 | BD Bioscience | 25x |
| **BB790** | CD161 | 624296 | DX12 | BD Bioscience | 25x |
| **PECy5** | CD28 | 555730 | CD28.2 | BD Bioscience | 10x |
| **PECy7** | CD25 | 335789 | 2A3 | BD Bioscience | 50x |
| **PECF594** | CD11c | 562393 | B-ly6 | BD Bioscience | 50x |
| **APC** | CD123 | 560087 | 7G3 | BD Bioscience | 25x |
| **APC-Cy7** | CD45 | 560178 | 2D1 | BD Bioscience | 200x |
| **BV421** | CD57 | 563896 | NK-1 | BD Bioscience | 400x |
| **BV510** | CD86 | 305432 | IT2.2 | Biolegend | 25x |
| **BV570** | CD3 | 624298 | UCHT1 | BD Bioscience | 100x |
| **BV605** | PD-1/CD279 | 329924 | EH12.2H7 | Biolegend | 25x |
| **BV650** | CD27 | 302828 | O323 | Biolegend | 100x |
| **BV711** | CD14 | 301838 | M5E2 | Biolegend | 100x |
| **BV750** | TIM3/CD366 | 624380 | 7D3 | BD Bioscience | 100x |
| **BV786** | CD127 | 351330 | A019D5 | Biolegend | 25x |
| **BUV563** | CD8 | 565695 | RPA-T8 | BD Bioscience | 200x |
| **BUV563** | CD19 | 741361 | HIB19 | BD Bioscience | 100x |
| **BUV615** | CD4 | 624297 | RPA-T4 | BD Bioscience | 400x |
| **BUV615** | IgD | 624297 | IA6-2 | BD Bioscience | 400x |
| **BUV661** | HLA-DR | <u>612980</u> | G46-6 | BD Bioscience | 100x |
| **BUV737** | CD45RO | 748368 | UCHL1 | BD Bioscience | 400x |
| **BUV805** | CD16 | 624287 | 3G8 | BD Bioscience | 400x |
| **AF700** | CCR7 | 353244 | G043-H7 | Biolegend | 25x |

#### Statistical analysis

Statistical significance was determined by GraphPad Prism using methods stated in the figure legends. P≤ 0.05 was considered significant. Details of the statistical tests are provided in each of the figure legends.

## Acknowledgements

This work was supported by Singapore National Medical Research Council’s grants NMRC/CIRG/1351/2013, NMRC/CSA-SI/0016/2017, NMRC/CIRG/1479/2017 awarded to S.G.L; NMRC/TCR/14-NUHS/2015, and NMRC/OFLCG19May-0038 grants awarded to J.E.C, G.P, S.G.L, R.D. This work was supported by CZI Seed Network grant CZIF2019-002429 awarded to R.D. We would like to thank Prof. Dr. Antonio Bertoletti and Dr. Nina Le Bert, Duke-NUS Medical School, for their critical feedback in improving the manuscript. We would like to thank Dr. Hwee Kuan Lee, BII, A*STAR for his support and guidance. We would like to thank Mr. Samydurai Sudhagar and the Genome Institute of Singapore’s S2GP facility (Spatial and Single Cell Platform) for the support. We thank Ms Ming Jie Lim in her assistance to the scRNAseq experiments. We thank Mr. Weiyong Chua for his administrative support. We thank Ms Xiaoning Wang and NUSMed Flow Cytometry Laboratory Unit for their support.

