## Supplementary Figure 1 for "Single cell profiling of functionally cured Chronic Hepatitis B patients reveals the emergence of activated innate and an altered adaptive immune response in the intra-hepatic environment"

**Supplementary Figure S1:**

Dot plot depicting the expression of key genes in each of the T cell subtypes

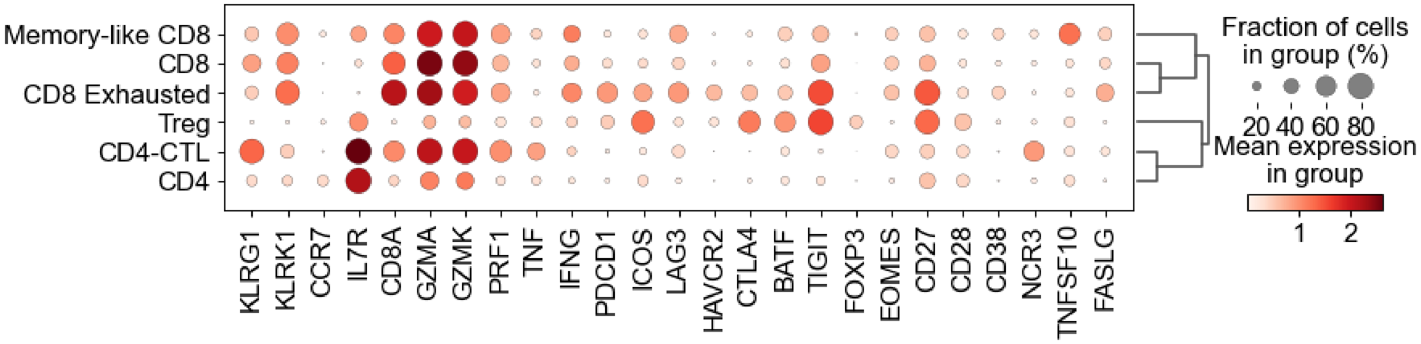
